# Lmx1a drives Cux2 expression in the cortical hem through activation of a conserved intronic enhancer

**DOI:** 10.1101/368555

**Authors:** Santiago P. Fregoso, Brett E. Dwyer, Santos J. Franco

## Abstract

During neocortical development, neurons are produced by a diverse pool of neural progenitors. A subset of progenitors express the *Cux2* gene and are fate-restricted to produce certain neuronal subtypes, but the upstream pathways that specify these progenitor fates remain unknown. To uncover the transcriptional networks that regulate *Cux2* expression in the forebrain, we characterized a conserved *Cux2* enhancer that we find recapitulates *Cux2* expression specifically in the cortical hem. Using a bioinformatic approach, we found several potential transcription factor (TF) binding sites for cortical hem-patterning TFs. We found that the homeobox transcription factor, Lmx1a, can activate the *Cux2* enhancer *in vitro*. Furthermore, we show that multiple Lmx1a binding sites required for enhancer activity in the cortical hem *in vivo*. Mis-expression of Lmx1a in neocortical progenitors caused an increase in *Cux2*^+^-lineage cells. Finally, we compared several conserved human enhancers with cortical hem-restricted activity and found that recurrent Lmx1a binding sites are a top shared feature. Uncovering the network of TFs involved in regulating *Cux2* expression will increase our understanding of the mechanisms pivotal in establishing *Cux2*-lineage fates in the developing forebrain.

**Summary Statement:** Analysis of a cortical hem-specific *Cux2* enhancer reveals role for *Lmx1a* as a critical upstream regulator of *Cux2* expression patterns in neural progenitors during early forebrain development.

## Introduction

During forebrain development, neural progenitor cells give rise to many different types of neuronal and glial cells that form the various telencephalic structures and circuits. This vast cellular diversity arises through the interplay between early tissue patterning pathways and gene regulatory networks (GRNs). Early in development, multiple tissue organizers and signaling centers provide morphogenic cues that govern regional identity and size. Within these different regions, complex transcriptional programs further diversify multipotent progenitor cells toward specific cell fates. The transcription factors (TFs) that establish the different GRNs to specify cell fates often work by binding gene regulatory elements, such as enhancers, to boost or suppress expression of target genes. Following transcriptional activation of GRNs, neural progenitors divide and eventually differentiate into specified cells. A key to better understanding forebrain development and function is to identify the signaling and transcriptional networks that establish regional identity and subtype fate specification during embryonic development.

The TF Cut-like homeobox 2 (Cux2) is dynamically expressed in complex spatiotemporal patterns in the developing mouse forebrain (Zimmer et al., 2004). During early brain development, a subset of neural progenitors weakly express *Cux2* transcripts in a salt and pepper pattern (Franco et al., 2012). We previously fate-mapped the lineage output of *Cux2*^+^ progenitors in the neocortex and found that this subset of neural progenitors are fate-restricted to produce late-born corticocortical neurons in upper layers (Franco et al., 2012; Gil-Sanz et al., 2015). Our studies indicated that *Cux2*^+^ progenitors in the developing forebrain are committed to this fate even before the onset of neurogenesis. However, the underlying mechanisms that restrict *Cux2*^+^ progenitors to specific cell fates remain largely unknown. *Cux2* knockout mice do not display any significant phenotype with respect to progenitor cell fate specification (Cubelos et al., 2008), implying that Cux2, while a useful marker for a fate-committed progenitor population, does not necessarily instruct fate in this context. We reasoned that a deeper understanding of Cux2^+^ cell fate commitment in forebrain progenitors could be achieved by uncovering the upstream GRNs responsible for the complex patterns of *Cux2* expression. Interestingly, neural progenitors in the dorsal telencephalic midline (DTM) strongly express *Cux2* in a more complete pattern than progenitors in adjacent regions, suggesting that this forebrain region might contain critical transcriptional regulators of the *Cux2* locus.

Previous studies have uncovered enhancers active in the developing mouse telencephalon, including an 856 bp element in intron 2 of the *Cux2* genomic locus that could drive strong transgene expression in the DTM (Hasenpusch-Theil et al., 2012; Visel et al., 2008). Here, we characterized this element as an active enhancer in the developing forebrain and show that it is specifically active in the cortical hem, but not in the adjacent hippocampus or neocortex. We further analyzed this enhancer for possible upstream regulators of *Cux2* expression. Among several bioinformatically identified candidates, we tested several transcription factors known to function in or be expressed within the cortical hem. Using an *in vitro* approach, we demonstrate that Lmx1a is a strong activator of the *Cux2* hem-specific enhancer. Additionally, *in vivo* Lmx1a gain-of-function in the neocortex, a region normally devoid of Lmx1a expression, increased the proportion of *Cux2*^+^ cells. Finally, we analyzed other enhancers that exhibit specific activity in the cortical hem and identify recurrent Lmx1a binding sites as a common motif shared between these distinct hem-specific enhancers. Our results suggest Lmx1a functions as an upstream regulator of a conserved *Cux2* enhancer in the cortical hem, and raise the possibility that Lmx1a is a critical TF in the GRN that specifies cortical hem fate.

## Results

### Early forebrain expression of *Cux2* begins at the dorsal telencephalic midline

To better understand when and where the earliest transcriptional regulators of *Cux2* are active in the developing telencephalon, we sought to define the temporal and spatial patterns of *Cux2* gene expression. We crossed *Cux2^Cre/+^* mice to the *Ai9* Cre-reporter line and used recombination (tdTomato^+^) as a readout of the cumulative transcriptional history of the *Cux2* genomic locus. *Cux2^Cre^/+;Ai9^fl/+^* brains were analyzed at E9.5, 10.5, 12.5 and 14.5 (Fig 1). We found that the earliest consistent pattern of recombined cells in the forebrain first appeared in the dorsal telencephalic midline (DTM) at ∼ E9.5 (Fig. 1A). At this age, a few recombined cells also began to appear scattered very sparsely throughout the adjacent neocortical neuroepithelium (Fig. 1A). By E10.5, the entire DTM was recombined, and the number of tdTomato^+^ neuroepithelial cells was increased in the neocortex (Fig. 1B). At E12.5 and E14.5, the DTM is reorganized to comprise 2 distinct structures: the cortical hem and choroid plexus epithelium (Grove et al., 1998). Essentially all cells in the cortical hem and choroid plexus were recombined at E12.5 (Fig. 1C) and E14.5 (Fig. 1D). In contrast, only a fraction of cells in the adjacent hippocampal primordium and neocortex were recombined (Fig. 1C-D). In fact, we observed a strikingly sharp border of complete-to-sparse recombination at the boundary between the cortical hem and the hippocampal primordium. These data indicate that forebrain activation of the *Cux2* locus occurs earliest and most uniformly in the DTM, including the cortical hem and choroid plexus.

**Fig. 1.**
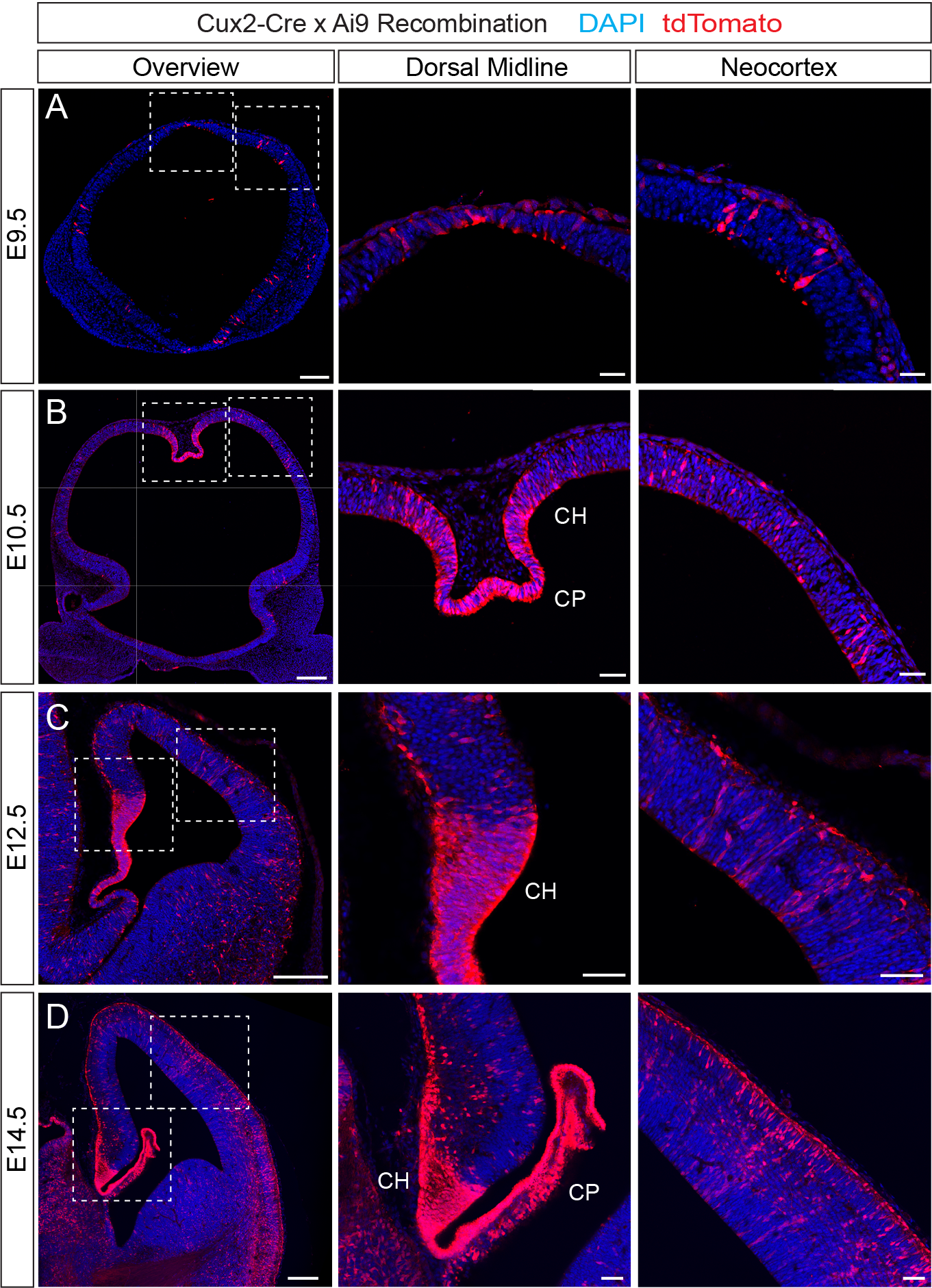
Spatiotemporal development of *Cux2* expression in murine telencephalic progenitors. Coronal sections of forebrains from *Cux2-Cre;Ai9* embryos showing recombination as a cumulative readout of *Cux2* expression. Recombined cells express tdTomato (red). Sections were counterstained for nuclei with DAPI (blue). Boxed insets show zoomed images of the dosral midline (middle panels) or neocortex (right panels). (A) E9.5: recombination is apparent in the dorsal-most region of the telencephalic neural tube and in scattered cells in the telencephalon. (B) E10.5: recombination becomes robust in the nascent cortical hem and choroid plexus, with salt-and-pepper recombination in the neocortex. (C) E12.5: recombination is nearly ubiquitous in the cortical hem, while still mosaic in the neocortex. (D) E14.5: recombination is complete in the cortical hem and much of the choroid plexus, while the neocortex exhibits a still expanding, mosaic pattern. Scale bars: left panels, 200 µm; middle and right panels, 50 µm, CH: cortical hem, CP: choroid plexus.

### *Cux2* regulatory element contains characteristics of an active enhancer and recapitulates the endogenous *Cux2* expression pattern in the cortical hem

Non-coding gene regulatory elements, such as enhancers, can act as critical platforms for TFs that drive cell fate decisions (Pattabiraman et al., 2014). To gain insights into some of the transcriptional programs that specify area and subtype fate in the telencephalon, we sought to identify enhancers that could recapitulate *Cux2* expression in the developing forebrain. A previous study identified an 856 base pair (bp) region within the human *Cux2* gene (hs611) that exhibits extreme human-rodent sequence conservation, suggesting an important functional role for this non-coding element (Visel et al., 2008). Indeed, both the human element (Visel et al., 2008) and the corresponding murine region (Hasenpusch-Theil et al., 2012) can drive restricted expression of a *lacZ* reporter gene in transgenic mouse embryos, indicating their role as functional enhancer elements. This enhancer lies within intron 2 of the *Cux2* gene (Fig. 2A) and has characteristics of an active enhancer in E14.5 forebrain tissue, including a prominent DNaseI hypersensitivity peak and histone marks H3K4me1 and H3K27ac, indicative of open, transcriptionally active chromatin (Fig. 2B).

**Fig. 2.**
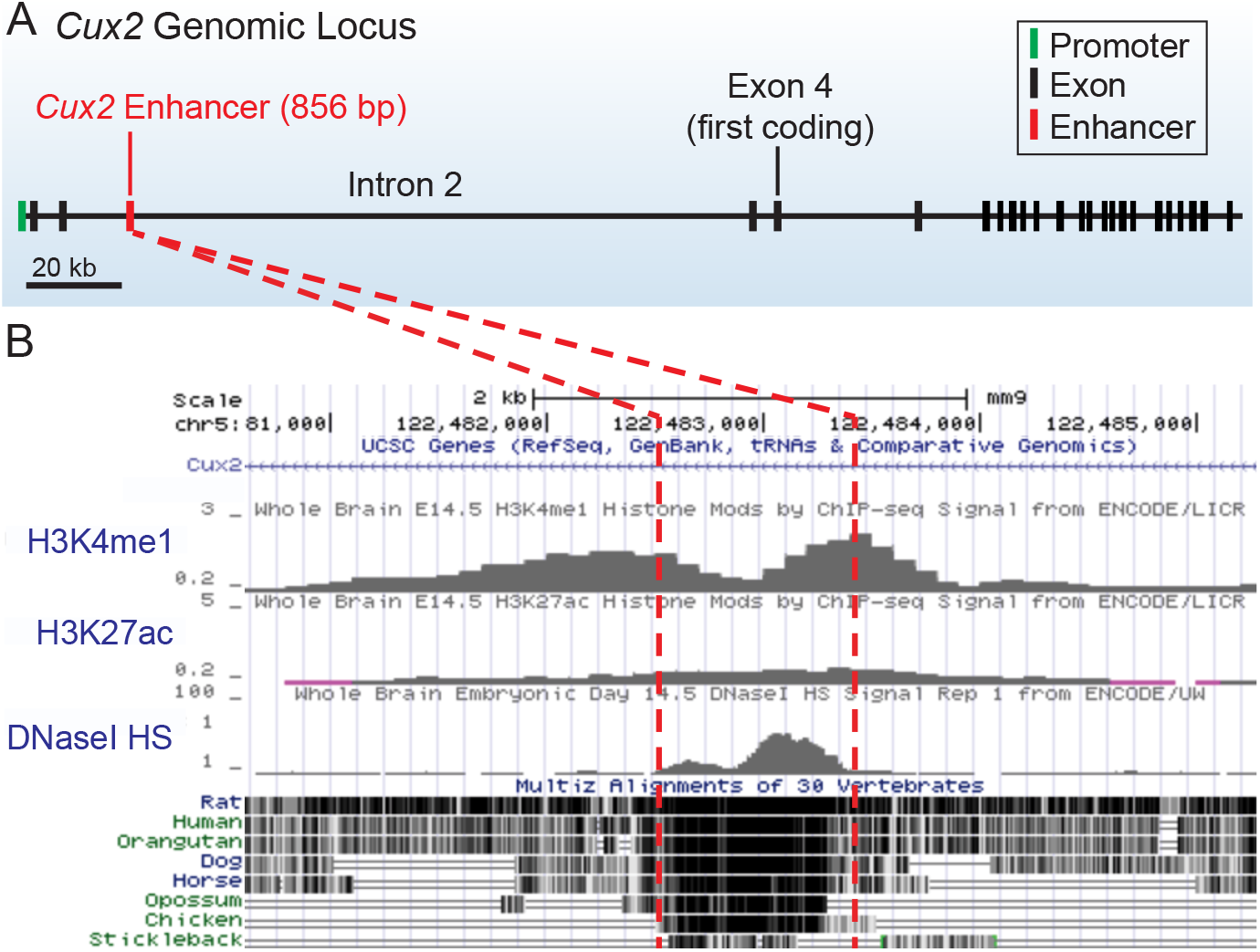
Genomic location and chromatin characteristics of a *Cux2* enhancer. (A) Schematic of murine *Cux2* genomic locus, showing the location of an 856 bp cis-regulatory element in the proximal region of intron 2. (B) UCSC Genome Browser data demonstrating key enhancer characteristics for the *Cux2* 856 bp element, including epigenetic marks H3K4me1 and H3K27ac, a prominent DNaseI hypersensitivity peak, and a high degree of evolutionary conservation across taxa.

Interestingly, the human and murine elements both exhibited expression patterns in the developing forebrain similar to that of *Cux2*, including strong expression in the DTM (Hasenpusch-Theil et al., 2012; Visel et al., 2008). To better characterize the expression pattern of this candidate enhancer in the developing forebrain, we first cloned the 856 bp murine region into an expression vector (Wilken et al., 2015) with a minimal promoter (TATA box) driving Cre recombinase. We then introduced the plasmid into the developing forebrain of *Ai9* Cre-reporter mice at E12.5, using *in utero* electroporation (Fig. 3A). We co-electroporated a plasmid expressing GFP from the ubiquitously-expressed synthetic CAG promoter (Niwa et al., 1991) as a marker of electroporated cells (Fig. 3A). Electroporations were performed to target different regions of the telencephalon, including the cortical hem, hippocampal primordium, and neocortex. We analyzed patterns of GFP expression and Cre-mediated recombination (tdTomato expression) at E14.5. As controls, we compared recombination patterns in the *Cux2*Enhancer-Cre electroporations to those of the minimal promoter construct alone (MINp-Cre, no enhancer) or with a strong and ubiquitous promoter (CAG-Cre). We found that recombined tdTomato^+^ cells in the MINp-Cre electroporations were very sparse in the DTM (Fig. 3B), hippocampus and the neocortex (Fig. 3E), consistent with weak expression from the TATA box alone. Conversely, the CAG-Cre construct drove recombination ubiquitously throughout the electroporated regions, including in the cortical hem, hippocampus and neocortex (Fig. 3C,F). Interestingly, recombination in the *Cux2*Enhancer-Cre electroporations was almost completely restricted to the cortical hem (Fig. 3D). Very few tdTomato^+^ cells were present in the hippocampus (Fig. 3D) or the neocortex (Fig. 3G). These data demonstrate that the activity of this *Cux2* enhancer is restricted within the telencephalon to the cortical hem, recapitulating a specific aspect of the complex endogenous *Cux2* expression pattern.

**Fig. 3.**
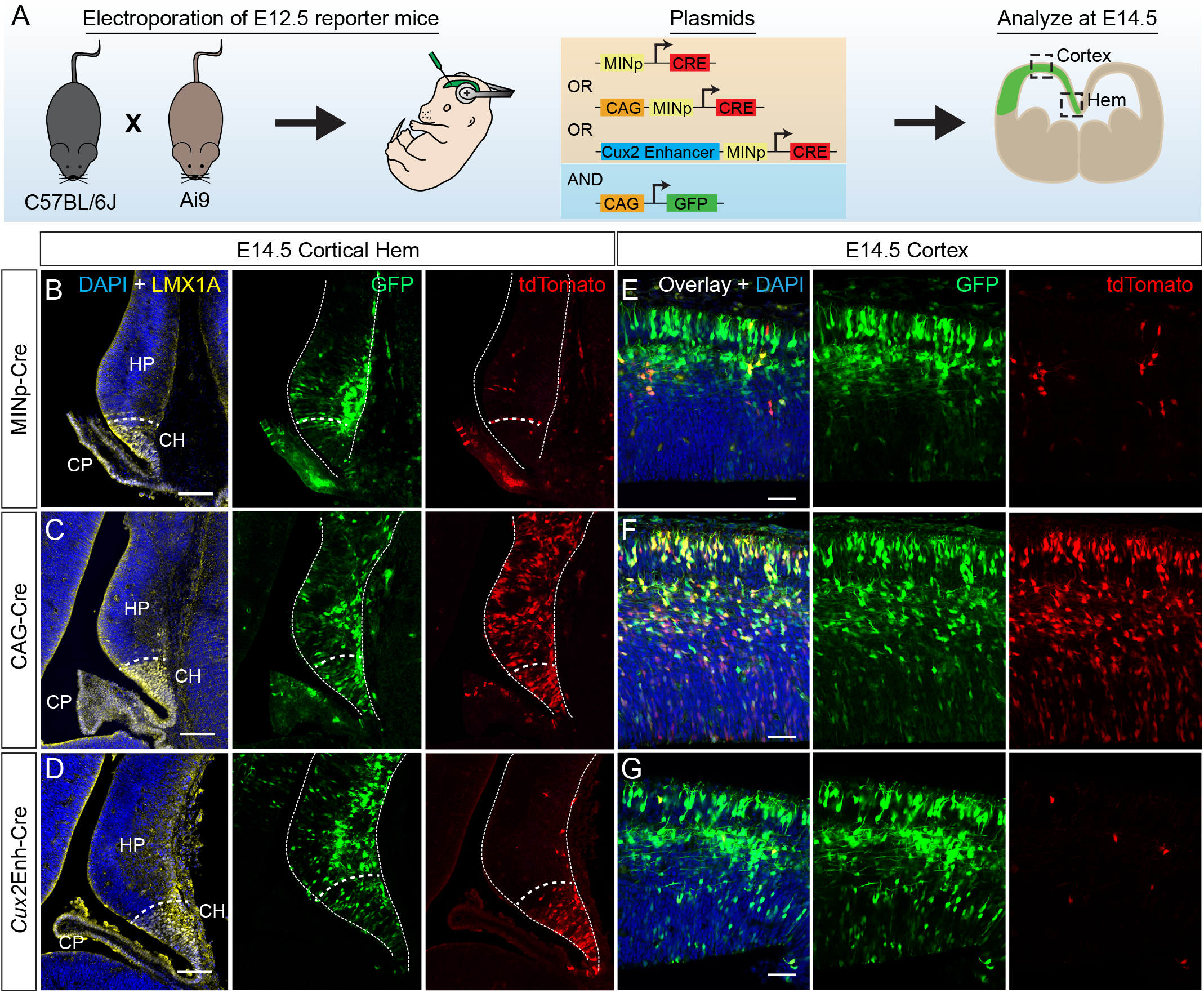
*Cux2* enhancer exhibits activity restricted to the cortical hem. (A) Schematic of experimental workflow: E12.5 *Ai9* reporter forebrains were electroporated *in utero* with constructs expressing Cre recombinase driven by either a minimal TATA-box promoter (Minp), ubiquitous CAG promoter, or the *Cux2* enhancer. CAG-GFP was co-electroporated to mark electroporated cells. Forebrains were harvested at E14.5 for analysis. (B-D) Coronal sections of electroporated brains showing the dorsal midline region. All electroporated cells express GFP (green) and recombined cells express tdTomato (red). Boundary between the cortical hem and hippocampal primoridum is marked by expression of LMX1A protein (yellow). Sections were counterstained for nuclei with DAPI (blue). Electroporation of Minp-Cre causes minimal recombination in the cortical hem and hippocampal primordium (B), whereas CAG-Cre leads to ubiquitous recombination throughout the electroporated regions (C). *Cux2*Enhancer-Cre drives robust recombination in the cortical hem, but not in the adjacent hippocampal primordium. (E-G) Coronal sections of electroporated brains showing the neocortex. Similar to the dorsal midline, Minp-Cre drives minimal recombination (E) and CAG-Cre drives ubiquitous recombination (F) in the neocortex. The *Cux2* enhancer (G) exhibits no activity in the neocortex. Scale bars: B-D, 100 µm; E-G, 50 µm. CH, cortical hem; CP, choroid plexus; HP, hippocampal primordium.

### Developmentally expressed cortical hem transcription factors, Lmx1a and Emx2, activate the *Cux2* enhancer

As pioneering regulators of development, TFs often control gene expression by acting on enhancers. To identify candidate transcriptional regulators of *Cux2* expression in the cortical hem, we analyzed the hem-specific enhancer sequence for predicted TF binding sites using the JASPAR (Khan et al., 2018) database with a ‘predicted’ and ‘consensus’ match threshold of 80% or higher. We identified ten recurring 8 base-pair clusters that contained consensus binding sequences for a set of known cortical hem-expressed TFs, including Emx2, Lmx1a and Msx1 (Fig. 4A-B). Lmx1a and Msx1 are expressed in the DTM (Fig. 4B) at E8.5 and E9.5, respectively, and their expression continues into adulthood (Failli et al., 2002; Furuta et al., 1997). Emx2 is expressed by neural progenitors in the hippocampus and neocortex beginning at E8.5, with the dorsomedial-most expression domain extending into the cortical hem (Fig. 4B) (Simeone et al., 1992a; Simeone et al., 1992b; Yoshida et al., 1997). As key players in telencephalic patterning, these TFs serve as ideal candidates for regulating *Cux2* expression in the cortical hem during early forebrain development. To test whether any of these TFs can activate the hem-specific *Cux2* enhancer, the enhancer element was cloned into the minimal promoter vector driving nuclear mCherry expression (Fig 5A). cDNAs for the candidate TFs were cloned into the bicistronic expression vector pCIG (Hand et al., 2005), which drives expression of both the TF and GFP from the CAG promoter. Each candidate TF plasmid was co-transfected with the *Cux2-* Enhancer-mCherry plasmid into murine immortalized neuroectodermal (NE-4C) cells (Schlett and Madarász, 1997). 24 hours after transfection, mRNA was collected from the cells and analyzed by RT-qPCR for levels of the *mCherry* reporter transcript (Fig. 5A). Compared to the pCIG empty vector negative control, Emx2 and Lmx1a significantly upregulated expression from the *Cux2* enhancer (Fig. 5B). In contrast, Msx1 did not change *Cux2* enhancer-driven *mCherry* levels compared to control (Fig. 5B).

**Fig. 4.**
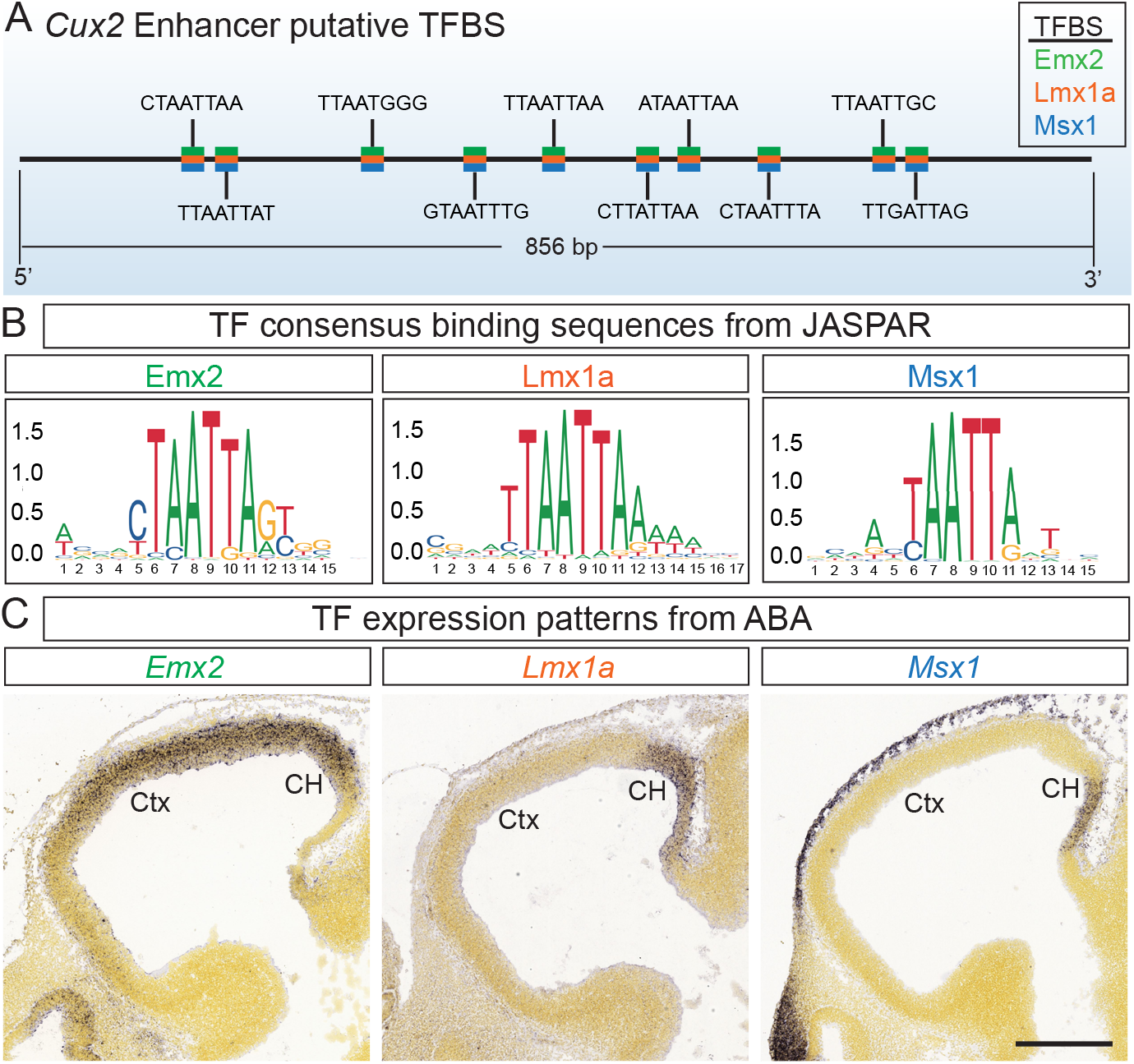
The *Cux2* enhancer contains multiple predicted binding sites for forebrain-patterning transcription factors expressed in the cortical hem. (A) Schematic of the *Cux2* enhancer with putative binding sites for Emx2, Lmx1a and Msx1, predicted from the JASPAR database at >80% threshold. (B) JASPAR motifs for consensus binding site sequences of the candidate TFs. (C) Sagittal sections from the Allen Brain Atlas *in situ* hybridization database showing mRNA expression of candidate TFs in the cortical hem of E11.5 mouse forebrains. Scale bar, 400 µm. Ctx, neocortex; CH, cortical hem.

**Fig. 5.**
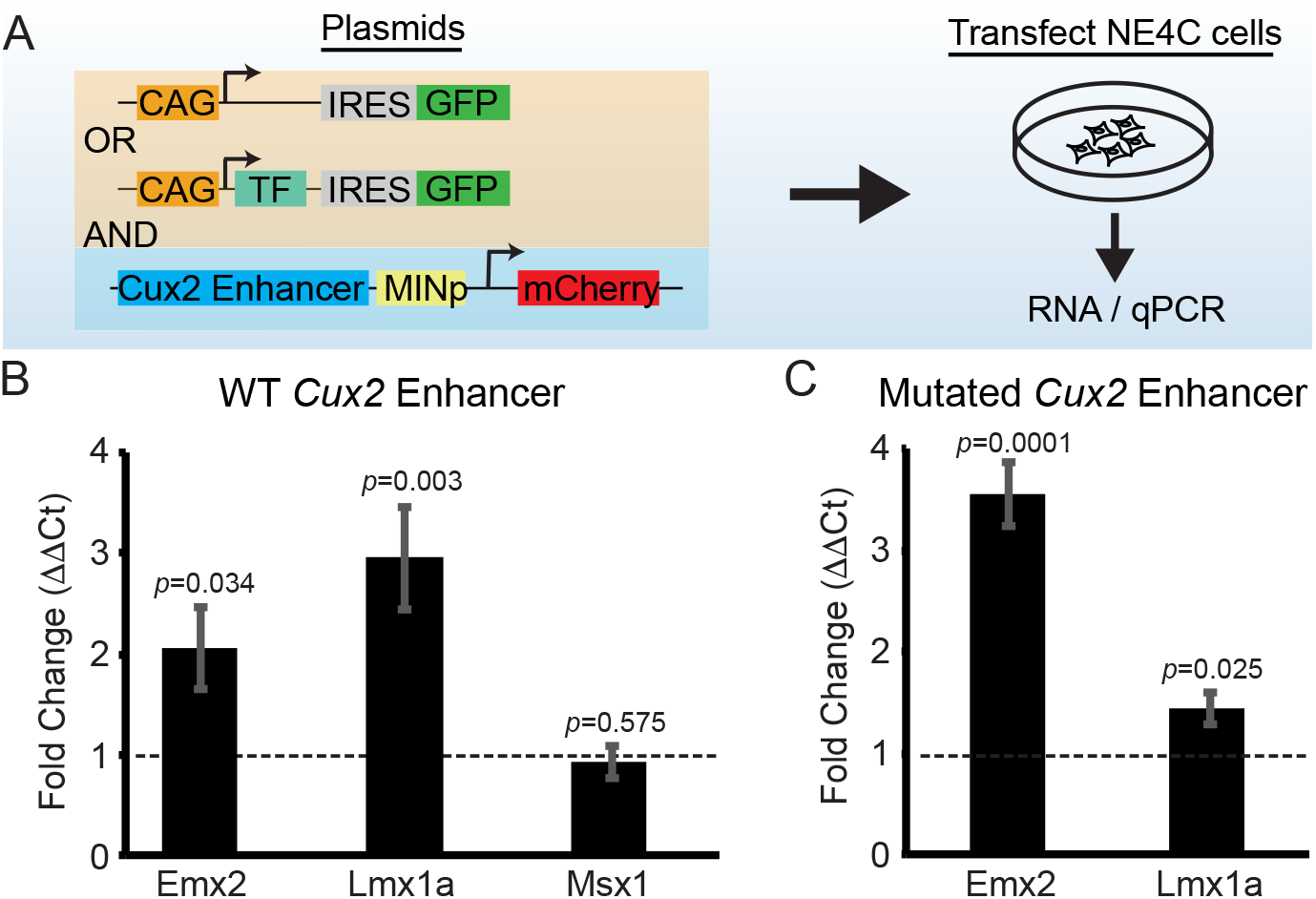
Lmx1a strongly activates the *Cux2* enhancer *in vitro*. (A) Schematic of experimental workflow. The *Cux2*Enhancer-mCherry plasmid was transfected into NE-4C cells together with either empty pCIG vector (CAG-IRES-GFP) or pCIG that expresses candidate TFs (CAG-TF-IRES-GFP). The effects of candidate TFs on expression of *Cux2*Enhancer-mCherry in NE-4C cells was quantified by qPCR of *mCherry* mRNA. (B-C) qPCR quantification of *mCherry* transcripts. Bar graphs are fold change (± SEM) over pCIG vector alone (dotted line), using the DDCt method. (B) Expression from the Cux2 enhancer is activated by expression of Emx2 and Lmx1a, but not Msx1. (C) When putative TF binding sites were mutated in the *Cux2* enhancer, Emx2 could still activate transcription but the effects of Lmx1a on transcription were greatly diminished.

### Activation of the *Cux2* enhancer in the cortical hem requires Lmx1a binding sites

Our *in silico* data predicted multiple binding sites for Emx2 and Lmx1a that overlapped each other (Fig. 4B), which correlated well with our *in vitro* studies demonstrating that these factors can activate the *Cux2* enhancer (Fig. 5B). To directly test whether these putative TF binding sequences were required for enhancer activation, we generated a mutant enhancer construct in which the central 8 base pairs of the predicted binding sites were mutated (Fig. 5C) and the mutant *Cux2*Enhancer-mCherry plasmid was tested for activation by Emx2 and Lmx1a. In contrast to the wild-type *Cux2* enhancer, the mutated version was no longer activated by Lmx1a (Fig. 5C). Surprisingly, Emx2 was still able to upregulate expression from the mutated enhancer (Fig. 5C), raising the possibility of other more critical Emx2 binding sites within the enhancer.

We next tested the activity of the mutated enhancer *in vivo* by *in utero* electroporation of the mutated enhancer driving Cre recombinase into *Ai9* reporter embryos (Fig. 6A). As we observed previously, the wild-type *Cux2*-Enhancer-Cre drove recombination specifically and robustly in the cortical hem (Fig. 6B). In contrast, the TF binding site mutant *Cux2*-Enhancer-Cre construct was unable to drive any recombination at all in the cortical hem (Fig. 6C). Together with our *in vitro* studies, these *in vivo* data indicate that the binding sites for Lmx1a are required for activity of the *Cux2* enhancer and that Lmx1a is an important transcriptional regulator of the *Cux2* enhancer in the cortical hem. The fact that mutating the enhancer abolished expression *in vivo*, but not activation by Emx2 *in vitro*, indicates that Emx2 may not be a critical regulator of the *Cux2* hem enhancer *in vivo*.

**Fig. 6.**
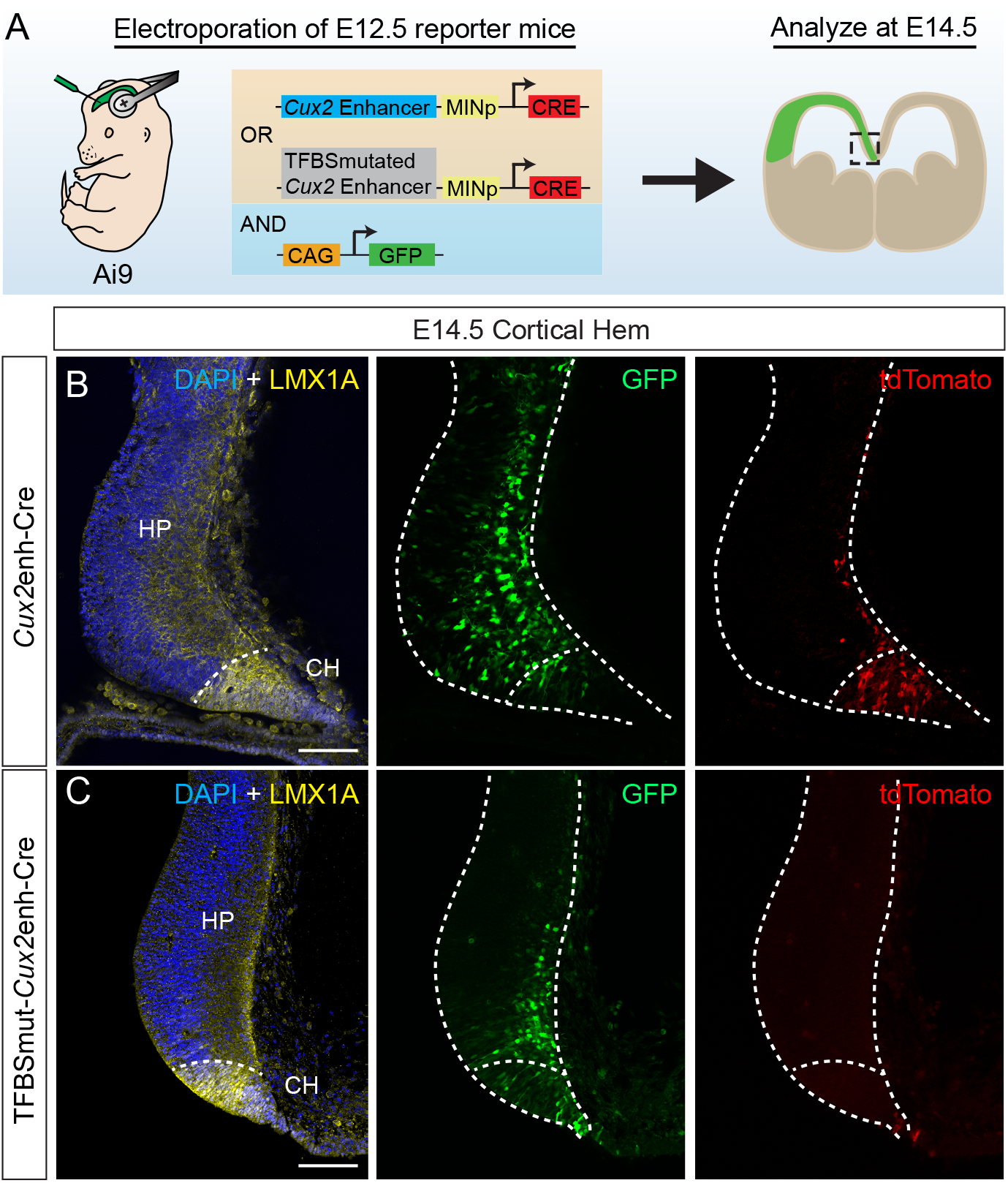
TFBS-mutated *Cux2* enhancer activity abolished *in vivo*. (A) Schematic of experimental workflow. *Ai9* reporter embryos were electroporated *in utero* at E12.5 with either the wild-type *Cux2* enhancer-Cre plasmid or the mutated version that is no longer activated by Lmx1a. CAG-GFP was co-electroporated as a marker of the electroporated cells. Forebrains were harvested at E14.5 for analysis of recombination. (B-C) Coronal sections of electroporated brains showing the dorsal midline region. All electroporated cells express GFP (green) and recombined cells express tdTomato (red). Boundary between the cortical hem and hippocampal primoridum is marked by expression of LMX1A protein (yellow). Sections were counterstained for nuclei with DAPI (blue). The wild-type *Cux2* enhancer driving Cre led to robust recombination specifically in the cortical hem (B), whereas the TFBS-mutated enhancer lost all activity in the cortical hem (C). Scale bars, 100 µm. Abbreviations as in Fig. 3.

### Lmx1a gain of function increases endogenous *Cux2* expression in the neocortex

In the developing telencephalon, Lmx1a is expressed strongly throughout the DTM, where it functions to promote cortical hem fate and suppress hippocampal and neocortical fate (Caronia-Brown et al., 2014; Chizhikov et al., 2010; Failli et al., 2002). The sharp border of *Lmx1a* expression between the cortical hem and hippocampus is very similar to that of *Cux2* (Fig. 7A), which is expressed strongly throughout the cortical hem but only weakly in a limited number of progenitors and neurons in the hippocampus and neocortex (Fig. 1). Since Lmx1a is not expressed at all in the developing neocortex where Cux2 expression is initially weak, this provided us an opportunity to assess whether mis-expression of Lmx1a in the neocortex is sufficient to upregulate endogenous *Cux2* expression. To test this possibility, we electroporated our CAG-Lmx1a-IRES-GFP construct into the neocortex of *Cux2*^Cre/+^; *Ai9^fl/+^* embryos *in utero* at E12.5 (Fig. 7B). We allowed the embryos to continue developing until E14.5 and analyzed the percentage of electroporated cells (GFP^+^) that belonged to the Cux2 lineage (tdTomato^+^). Compared to the CAG-IRES-GFP control (Fig. 7C-D), Lmx1a mis-expression resulted in a statistically significant (*p* = 0.0001) 2-fold increase in tdTomato^+^ cells within the electroporated population (Fig. 7E-G). These data support a role for Lmx1a in promoting endogenous *Cux2* expression in the early developing telencephalon.

**Fig. 7.**
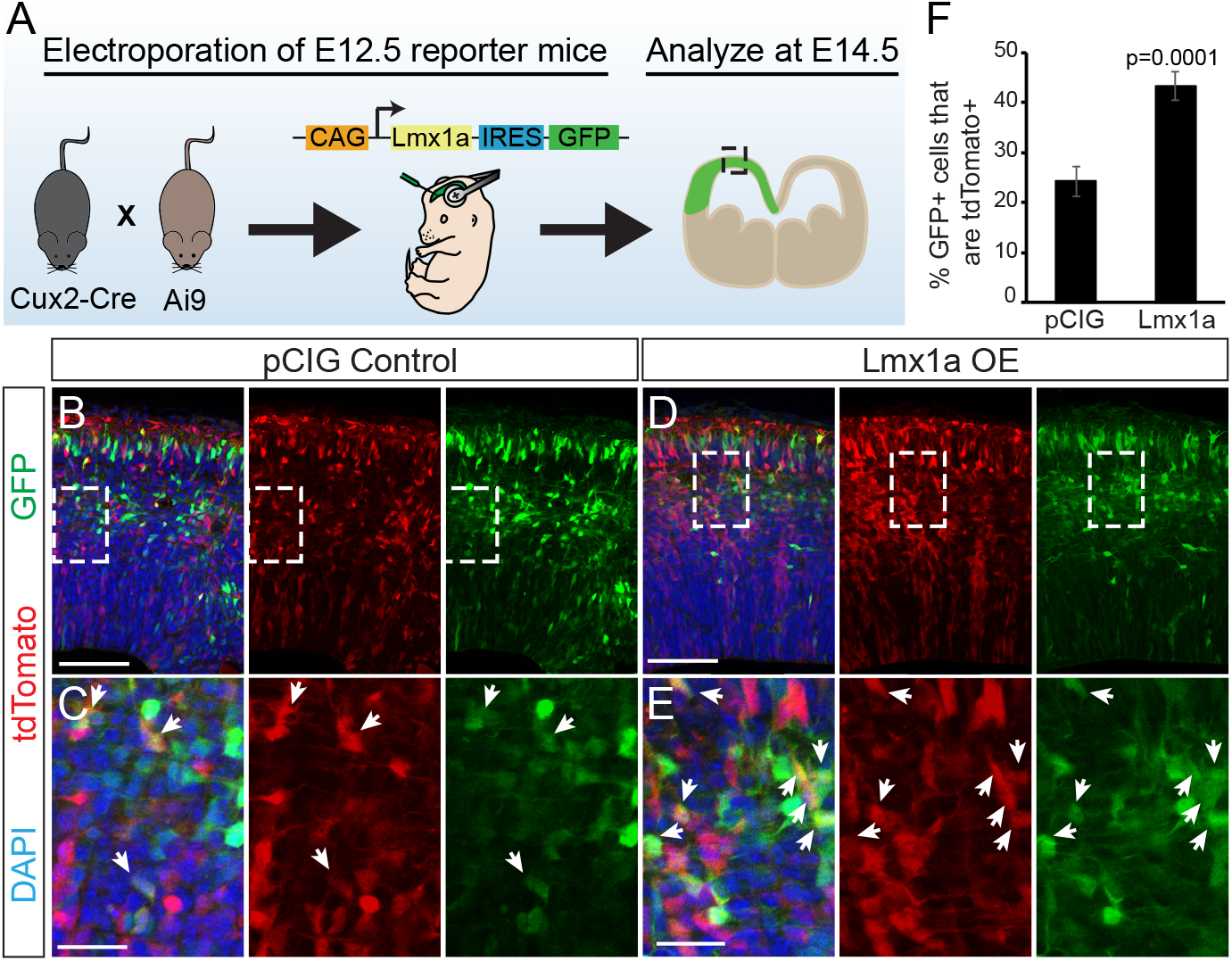
Lmx1a gain-of-function in the neocortex increases the number of *Cux2*^+^ cells. (A) Schematic of experimental workflow. *Cux2-Cre;Ai9* embryos were electroporated at E12.5 with a constitutive Lmx1a expression plasmid. Forebrains were harvested at E14.5 for quantification of the percentage of electroporated cells that were recombined. (B) Coronal section of a neocortex electroporated with empty vector control (pCIG). Electroporated cells are GFP^+^ (green), and recombined cells from the Cux2-Cre lineage are tdTomato^+^ (red). Sections were counterstained for nuclei with DAPI (blue). (C) Magnified inset from (B) showing electroporated *Cux2*-lineage cells. (D) Coronal section of a neocortex electroporated with the Lmx1a expression vector. GFP^+^ cells are mis-expressing Lmx1a. (E) Magnified inset from (D) showing that Lmx1a gain-of-function results in an increased percentage of electroporated cells that are recombined. (F) Quantification showing percent (± SEM) of electroporated cells (GFP^+^) that are recombined (tdTomato^+^) in control vs Lmx1a electroporations. Scale bars: B and D, 100 µm; C and E, 25 µm.

### Multiple Lmx1a binding sites are a shared feature among cis-regulatory elements active in the cortical hem

Previous work has revealed a number of forebrain enhancers, including some that appear to have restricted cortical hem activity similar to the murine *Cux2* enhancer (Pattabiraman et al., 2014; Visel et al., 2008). Human enhancer elements hs411, hs611 and hs643 can drive cortical hem-specific lacZ expression in transgenic mouse embryos (Fig 8A-C). We reasoned that all three enhancers might share features that regulate their activity through a common mechanism, given their very similar spatial and temporal transcriptional activity. To uncover common features between hs611, hs411 and hs643, the sequences of all three genomic regions together with the murine *Cux2* hem enhancer were analyzed using Analysis of Motif Enrichment (McLeay and Bailey, 2010) through the MEME Suite web portal (http://meme-suite.org/tools/ame). Compared to 1004 shuffled control sequences, the most enriched motif shared by all four elements was a TTAATTAA motif (*p* = 1.48e-6 by Fisher’s exact test) that was identified as an Lmx1a consensus binding motif by JASPAR, Jolma and Uniprobe databases (Fig. 8D). We next used the JASPAR database (>85% threshold) to search all three human enhancer elements for putative Lmx1a binding sites. Similar to the murine *Cux2* cortical hem enhancer, hs611, hs411 and hs642 are all predicated to contain multiple high-threshold Lmx1a binding sites (Fig. 8E). As a common feature among cortical hem enhancers, the presence of multiple Lmx1a binding sites may indicate that Lmx1a sits near the top of the GRN active in the developing cortical hem, perhaps as a pioneering TF.

**Fig. 8.**
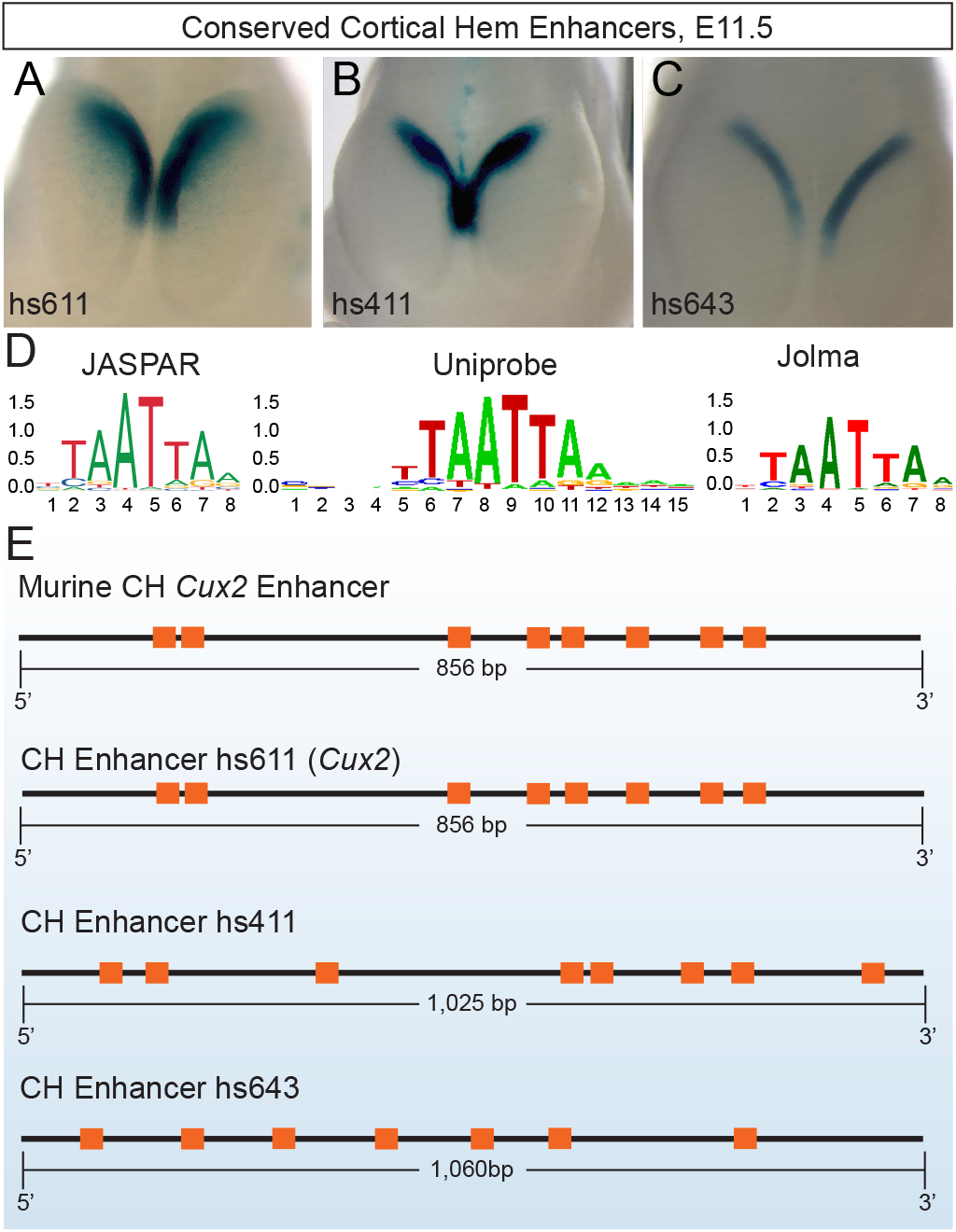
Lmx1a binding sites are a common feature of enhancers active in the developing cortical hem. (A-C) Examples of human enhancer elements driving LacZ expression in transgenic mouse embryos. Whole mount staining images from the Vista Enhancer Browser show activity of hs411 (A), hs611 (B) and hs643 (C) in the cortical hem region. (D) Analysis of Motif Enrichment identified Lmx1a consensus binding sites as the most significantly enriched motif in the 4 hem-expressed enhancers. (E) Using the JASPAR database, all four cortical hem enhancers were predicted to contain 7 or more high-threshold Lmx1a binding sites.

## Discussion

Using a combination of *in silico, in vitro* and *in vivo* approaches, we characterized a *Cux2* gene regulatory element with the goal of uncovering key TFs involved in the transcriptional regulation of *Cux2* in neural progenitors. We showed that this *Cux2* enhancer recapitulates a specific aspect of the complex *Cux2* expression pattern in the developing forebrain, namely strong and precise expression in the cortical hem. Our further analysis uncovered the LIM homeobox transcription factor, Lmx1a, as a positive regulator of the *Cux2* cortical hem enhancer. Comparison of 3 cortical hem-specific human enhancer elements revealed that recurring Lmx1a binding sites is the top shared motif, raising the possibility that Lmx1a is master transcriptional regulator of the GRN that controls cortical hem identity.

### *Cux2* expression as a tool to uncover regulators of cell-fate decisions

We previously identified a subset of neural progenitors in the developing forebrain that are fate-restricted to produce only corticocortical projection neurons in the neocortex (Franco and Müller, 2013; Franco et al., 2012; Gil-Sanz et al., 2015). These progenitors can be identified by expresssion of the transcription factor, *Cux2*, and lineage-traced using *Cux2-Cre* and *Cux2-CreERT2* knock-in mice. Although Cux2 itself does not appear to control cell fate decisions in the forebrain (Cubelos et al., 2008; Cubelos et al., 2010), its restricted expression in defined subsets of neural progenitors may be a useful tool for uncovering transcriptional regulators of cell fate during forebrain patterning. The *Cux2* expression pattern in the developing forebrain is complex and dynamic (Franco et al., 2012; Gil-Sanz et al., 2015; Zimmer et al., 2004), suggesting that control of the *Cux2* locus may involve multiple transcriptional regulatory mechanisms. Using our *Cux2-Cre* mice crossed to a Cre-reporter line, we identified the DTM as one of the earliest sites of *Cux2* expression in the developing forebrain. In contrast to the salt-and-pepper pattern of *Cux2* expression in the adjacent hippocampus and neocortex, we find that essentially all neural progenitors in the cortical hem belong to the *Cux2* lineage. This raised the interesting possiblity that *Cux2* expression in different parts of the developing forebrain are controlled by distinct mechanisms. Identification of the various cis-regulatory elements that drive differential *Cux2* expression, and the transcription factors that regulate these elements, may therefore lead to a better understanding of the GRNs that control tissue patterning and subtype fate specification in the developing forebrain.

### Identification and characterization of a cortical hem-specific *Cux2* enhancer

Previous studies identified genomic regions within intron 2 of the human and murine *Cux2* genes that can recapitulate *Cux2* expression in the DTM of transgenic mice (Hasenpusch-Theil et al., 2012; Visel et al., 2008). We found that the murine *Cux2* element exhibits features of an active enhancer in the E14.5 forebrain, including DNaseI hypersensitivity and histone modifications associated with transcriptionally active chromatin. This region also displays high levels of conservation from humans to chickens, pointing toward an important functional role. Using *in utero* electroporation to test the *in vivo* activity of this region, we show that it drives expression specifically and robustly in the cortical hem neuroepithelium, but not in progenitors in the adjacent hippocampus or neocortex. These data indicate that this element is a developmentally active enhancer specific for the cortical hem. It will be interesting in future studies to determine which features of this enhancer are required for *Cux2* expression in the cortical hem, and whether this regulatory element is active in other *Cux2* expression domains that share features with the cortical hem, such as the rhombic lip in the hindbrain (Capaldo and Iulianella, 2016).

### Lmx1a is a critical regulator of the *Cux2* hem-specific enhancer

Using a bioinformatics approach, we identified several putative TF binding sites in the *Cux2* enhancer. As known regulators of cortical hem development (Chizhikov et al., 2010; Tole et al., 2000), Emx2, 12 Lmx1a and Msx1 comprised a promising group of candidate TFs with the potential to regulate the *Cux2* enhancer in the cortical hem. We did not identify any effect of Msx1 on activity of the *Cux2* enhancer *in vitro*, indicating it may not be a direct regulator of *Cux2* cortical hem expression. On the other hand, both Emx2 and Lmx1a activated transcription from the *Cux2* enhancer. The majority of the predicted Emx2 and Lmx1a binding sites overlapped each other, reflective of the similarity of their consensus binding motifs. Interestingly, mutation of several of these putative binding sites drastically reduced responsiveness of the enhancer to Lmx1a, but not to Emx2. This may suggest substantial redundancy in the Emx2 binding sites in the *Cux2* enhancer, or that the remaining Emx2 binding sites are more critical. Importantly, the mutated enhancer showed no activity in the cortical hem, suggesting that binding of Lmx1a, but not Emx2, is critical for enhancer activation *in vivo*. This would be in line with our further experiments showing that the *Cux2* enhancer is not active in the neocortex, where Emx2 is strongly expressed but Lmx1a is absent.

In further support of Lmx1a as an activator of *Cux2* expression, we found that mis-expressing Lmx1a in the neocortex results in an increase in Cux2^+^ cells. Although this assay does not allow us to determine whether the increase results from direct activation of the cortical hem enhancer, our data point to Lmx1a as being sufficient to activate *Cux2* expression in the developing forebrain. Interestingly, Lmx1a mis-expression in the neocortex did not drive recombination in all cells in the electorporated region, in contrast to the complete recombination pattern seen in the cortical hem of *Cux2-Cre* mice. This result could simply be due to insufficent levels or duration of Lmx1a expression, or it may suggest the presence of additional factors that regulate *Cux2* expression. For example, Lmx1a may require a transcriptional co-activator for maximum activity that is missing from the neocortex, or perhaps there is an unidentified transcriptional repressor of the *Cux2* enhancer that is expressed specifically in the neocortex. Further studies will be required to fully elucidate the mechanisms that control the complex expression pattern of *Cux2* in the developing forebrain.

### Lmx1a as a common activator of cortical hem GRNs

When we compared the murine and human Cux2 enhancers to two other conserved human elements that drive expression in the cortical hem, we found that the top motif enriched in all four enhancers corresponds to the Lmx1a consensus binding site. Together with the fact that Lmx1a is one of the earliest markers of the DTM (Failli et al., 2002; Mangale et al., 2008), these data suggest that Lmx1a may sit near the top of the GRN involved in regulating cortical hem cell fate. In line with this idea, cortical hem identity is lost in *dreher* mutant mice in which *Lmx1a* is inactivated by a missense mutation (Chizhikov et al., 2010). The sharp border of Lmx1a expression between the cortical hem and adjacent hippocampal primordium further make it an ideal candidate for establishing precise patterns of gene expression during early patterning of the developing forebrain. An important unanswered question is what lies upstream of Lmx1a during these early patterning stages. Previous work has reported that *Lmx1a* expression can be activated by BMP4 in the developing forebrain (Srinivasan et al., 2014; Watanabe et al., 2016). As a morphogenetic pathway that is specifically expressed within the cortical hem and choroid plexus, BMP signaling could potentially initiate the Lmx1a-dependent GRN that leads to specific DTM fates. Interestingly, previous studies have reported that upregulation of BMP signaling both in the developing chick olfactory epithelium (Wittmann et al., 2014) and murine mandibular neural crest cells (Bonilla-Claudio et al., 2012) results in significant upregulation of *Cux2* expression. Additionally, *Cux2* expression appears coincident with BMP4 within the mesenchyme of the developing mouse limb bud (Iulianella et al., 2003). How BMPs activate *Cux2* expression in these contexts has not been determined, but it would be interesting to test whether BMP signaling can drive *Cux2* expression in multiple tissues through Lmx1a-mediated activation of the conserved enhancer.

## Conclusions

In this study we identify a conserved enhancer and its transcriptional activator, Lmx1a, as an important mechanism for driving restricted expression of *Cux2* in the developing forebrain. We further show that recurrent Lmx1a binding sites are a common motif shared in multiple enhancers with similarly restricted activities. These studies provide a template for future studies aimed at identifying other *Cux2* cis-regulatory elements that control its complex expression during forebrain development, and the ultimately the upstream GRNs that specify different cell fates among the forebrain progenitor pool.

## Materials and Methods

### Animals

Mice used for experiments were housed and handled in accordance with protocols approved by the UC Anschutz Medical Campus IACUC committee. The following mouse lines were used in this study: *Cux2-Cre* (Franco et al., 2012; 2011(Franco et al., 2012; Franco et al., 2011; Gil-Sanz et al., 2015)), *Ai9* (Madisen et al., 2010) and C57BL/6J. Embryos were produced from timed-pregnant females, with noon on the day of plug being designated E0.5.

### Plasmids and In Utero Electroporation

The murine *Cux2* enhancer was cloned from the endogenous genomic locus (NCBI37/mm9 chr5:122,482,512-122,483,367) using a Gblock (IDT) with 5’ and 3’ arms homologous to the multiple cloning site in the backbone vector. The Gblock was cloned by Gibson assembly into the pMinp vector (Wilken et al., 2015), immediately upstream of the TATA box. mCherry or Cre recombinase with a nuclear localization signal (Lewandoski and Martin, 1997) were cloned immediately downstream of the TATA box. To generate the TF binding site mutant version of the *Cux2* enhancer, we synthesized a Gblock in which the central 8 base pairs of each putative binding site was mutated to 5’ AAGCGCAA3’. Transcription factor cDNAs were either obtained from Addgene (Lmx1a: 45070, Msx1: 34998) or from IDT as Gene blocks (Emx2) and cloned into the Sac1 and Xma1 sites of the pCIG vector (Hand et al., 2005), between the CAG promoter and the IRES-GFP cassette. In utero electroporation of plasmids (0.5–1mg/ml) were carried out as previously described (Franco et al., 2012; Gil-Sanz et al., 2013) on E12.5 embryos of timed-pregnant mice. Embryos were harvested for analysis at E14.5.

### Immunohistochemistry

Brains from E9.5–14.5 embryos were dissected and fixed for 2 hours at room temperature in 4% paraformaldehyde. Forebrains were sectioned on a vibrating microtome (Leica VT1200S) at 100 µm increments, or on a cryostat (Leica CM1520) at 15–30 µm increments. Immunohistochemistry was performed on tissue sections as described previously (Winkler et al., 2018) using the following antibodies: rabbit anti-Lmx1a (1:1000, Millipore, RRID:AB_10805970), rabbit anti-RFP (1:500, LifeSpan Biosciences, RRID:AB_945213). Donkey secondary antibodies conjugated to Alexa Fluor 488, Rhodamine Red-X, or Alexa Fluor 647 were purchased from Jackson ImmunoResearch and used at 1:500. Sections were imaged using a Zeiss LSM 780 confocal microscope.

### Cell culture and qRT-PCR

Experiments were performed using the immortalized mouse neuroectodermal NE-4C cells (ATCC CRL-2925), grown in Dulbecco’s minimal essential media (MEM; Corning 10-010-CV) with 4mM L-glutamine (Invitrogen), 10% fetal bovine serum (FBS) (Invitrogen) and Penicillin (0.0637g/L)-Streptomycin (0.1g/L). Cells were plated on 12 well plates and grown to ∼70% confluency prior to transfection. Cells were transfected with either CAG-Emx2, Lmx1a or Msx1-IRES-GFP, together with *Cux2*Enhancer-mCherry or TF binding site mutated *Cux2*Enhancer-mCherry for 4–6 hrs with Lipofectamine 3000 (Invitrogen), with subsequent media change. 24 hrs following transfection, RNA was isolated from cells with an RNeasy Plus Kit (Qiagen) and reverse transcribed into cDNA using an iScript RT Kit (Bio-Rad). Expression of *mCherry, GFP*, and housekeeping gene *Cyclophilin A* was assessed by qRT-PCR (Bio-Rad CFX Connect R-T System). Fold change was calculated by the delta-CT method for both *GFP* and *mCherry*, relative to *Cyclophilin A*. Fold changes of *mCherry* mRNA were normalized to those of *GFP*, to account for variations in transfection efficiency. The following primers were used: *Cyclophilin A* forward: GAGCTGTTTGCAGACAAAGTTC, *Cyclophilin A* reverse: CCCTGGCACATGAATCCTGG, *eGFP* forward: ACGTAAACGGCCACAAGTTC, *eGFP* reverse: AAGTCGTGCTGCTTCATGTG, *mCherry* forward: GATAACATGGCCATCATCAAGGA, *mCherry* reverse: CGTGGCCGTTCACGGAG.

### Quantitative analysis of Lmx1a gain of function

CAG-Lmx1a was electroporated into cortices of E12.5 *Cux2^Cre/+^;Ai9^fl/+^* embryos (n=3) followed by quantification of *Cux2* expression at E14.5. At least 3 histological sections from distinct rostro-caudal regions collected from 3 different animals were analyzed in regions comprising primarily the somatosensory cortex. Single-plane confocal images were used for quantification. Cux2^+^ cells were counted based on tdTomato expression from the recombined Ai9 allele while Lmx1a expressing cells were labeled by GFP expression. *Cux2*-expressing cells were quantified as a percentage of those expressing Lmx1a. All analysis was performed using Fiji/ImageJ on 3–5 20x images per brain.

### Statistics

As all comparisons made were between two groups, a two-tailed, two-sample equal or unequal variance Student *t*-tests were used to analyze all data. Equality of variance was determined using a Bartlett’s Test. The standard error of the mean (SEM) is reported on all graphs.

## Acknowledgments

We thank Joseph Brzezinski (University of Colorado – Anschutz Medical Campus) for providing the minimal promoter plasmids.

## Competing interests

The authors declare no competing financial interests.

## Funding

This work was supported by the Children’s Hospital Colorado Program in Pediatric Stem Cell Biology and The Boettcher Foundation (S.J.F.).

